# PSIICOS projection optimality for EEG and MEG based functional coupling detection

**DOI:** 10.1101/2023.03.26.534253

**Authors:** Dmitrii Altukhov, Daria Kleeva, Alexei Ossadtchi

## Abstract

Functional connectivity is crucial for cognitive processes in the healthy brain and serves as a marker for a range of neuropathological conditions. Non-invasive exploration of functional coupling using temporally resolved techniques such as MEG allows for a unique opportunity of exploring this fundamental brain mechanism in a reasonably ecological setting.

The indirect nature of MEG measurements complicates the the estimation of functional coupling due to the spatial leakage effects. In previous work (Ossadtchi et al., 2018), we introduced PSIICOS, a method that for the first time allowed us to suppress the spatial leakage and yet retain information about functional networks whose nodes are coupled with close to zero or zero mutual phase lag.

In this paper, we demonstrate analytically that the PSIICOS projection is optimal in achieving a controllable trade-off between suppressing mutual spatial leakage and retaining information about zero-phase coupled networks. We also derive an alternative solution using the regularization-based inverse of the mutual spatial leakage matrix and show its equivalence to the original PSIICOS. This approach allows us to incorporate the PSIICOS solution into the conventional source estimation framework. Instead of sources, the unknowns are the elementary networks and their activation timeseries are formalized by the corresponding source-space cross-spectral coefficients.

Additionally, we outline potential avenues for future research to enhance functional coupling estimation and discuss alternative estimators that parallel the established source estimation approaches. Finally, we propose that the PSIICOS framework is well-suited for Bayesian techniques and offers a principled way to incorporate priors derived from structural connectivity.

## 1. Introduction

Functional connectivity, which refers to the interaction and communication between different brain regions, plays a crucial role in understanding various cognitive processes taking place in the brain (Greicius, 2008; Hutchison et al., 2013). Non-invasive techniques like magnetoencephalography (MEG) and electroencephalography (EEG) have greatly aided research in this field. These techniques provide high temporal resolution, which allows researchers to investigate various types of functional coupling between neuronal population activity time series. This includes identifying types of functional coupling that are marked by phase synchrony and distributed across different frequency bands (Siegel et al., 2012).

The cross-spectrum is frequently utilized to calculate functional connectivity between two narrow band neuronal timeseries (Gail et al., 2004; Zeitler et al., 2006; Bastos and Schoffelen, 2016). As shown in (Aydore et al., 2013) the absolute value of the normalized cross-spectrum measures the degree of phase coupling between a pair of circularly Gaussian random processes. Therefore, according to the *communication through coherence* principle (Fries, 2015; Varela et al., 2001), the coherence is often used to gauge the amount of communication between a pair of neuronal populations (Bowyer, 2016).

Investigating brain function using non-invasive, time-resolved imaging methods such as EEG and MEG offers access to a great level of details regarding the dynamics of neuronal activity combined with the versatility of the iterative experimental design. EEG and MEG represent indirect measurements of brain activity and yield multivariate timeseries where each channel represents a mixture of activity of individual neuronal sources. Such a mixing, known as spatial leakage (SL), hinders estimation of the cross-spectral coefficients between neuronal sources based on such non-invasively collected EEG and MEG. To partially resolve this problem, functional coupling estimation is typically preceded by a source timeseries estimation step using one of the numerous inverse solvers developed by the community over the last several decades (Samuelsson et al., 2021; Westner et al., 2022). The role of this step is to “unmix” the sensor data and gain access to the activation timeseries of the individual neuronal sources, approximated by equivalent current dipoles (ECDs). The inverse problem of EEG and MEG is fundamentally ill-posed, which means that no inverse method can achieve perfect unmixing. The amount of the remaining leakage from the other sources present in the estimate of activity of a given source depends on the inverse solver parameters (Hincapie et al., 2017) and significantly affects the accuracy of functional coupling estimates. The remaining SL still leads to inaccurate estimates of functional connectivity.

The use of the imaginary part of coherence protects against the SL effects (Nolte et al., 2004), but also reduces the sensitivity to the source pairs whose interaction is characterized by the timeseries with close-to-zero or zero mutual phase lags. This is because the coherence between a pair of signals with a short mutual time lag relative to the oscillation period predominantly shows up in the real part of the complex-valued cross-spectrum (Rabiner and Gold, 1975).

To address the issue of spatial leakage (SL) in functional connectivity estimation, we previously introduced a method called PSIICOS (Phase shift invariant imaging of coherent sources) in (Ossadtchi et al., 2018). PSIICOS operates in the space of vectorized cross-spectral matrices and employs a projection operation to minimize the contribution of the SL while retaining the information about zero or close-to-zero phase interactions. As demonstrated with realistic simulations and in application to real MEG data, the PSIICOS methodology augments more traditional approaches (Nolte et al., Stamet al., 2007; Nolte et al., 2008; Pascual-Marqui et al., 2011; Ewald et al., 2012) that rely solely on the imaginary part of the normalized cross-spectrum and therefore struggle in detecting functional networks with zero or close-to-zero phase coupling.

In this work we analytically show that PSIICOS is optimal in terms of minimization of the amount of global cross-talk for any pair of cortical locations tested for being the generators of functionally coupled timeseries. We start with formulating the appropriate optimality criterion and then analytically solve the associated optimization problem to arrive at the PSIICOS projection. We also derive an alternative solution based on the regularization principle and show its equivalence to the original PSIICOS. We then provide the results of the numerical analysis illustrating the ideas behind the optimal selection of the projection rank or regularization coefficient. We also assess the amount of benefit offered by operation in the *M*^2^ dimensional space by exploring the properties of lower rank approximation of PSIICOS filters. We conclude by discussing the potential of using *M*^2^ dimensional space operations to improve the estimation methods for functional coupling in MEG and EEG.

## 2. Spatial leakage and its suppression by means of orthogonal projection in the product space of sensor signals

### 2.1. Generative models of the sensor signals

The EEG/MEG experimental setup involves recording *K* epochs of electrophysiological activity using an *M*-channel encephalographic device. The subject’s brain anatomy, either from a generic head model or an individual MRI scan, defines the source space, a mesh of *N* vertices approximating cortical surface, so that each vertex harbors a neuronal source approximated by the equivalent current dipole (ECD). The *K* × *N* forward operator **G** resulting from Maxwell’s equations for electromagnetic activity maps the activity of these ECDs to the sensor-space measurements. Assuming for each of *K* trials identical configuration of active sources whose indices form set Ω the observed *k* − *th* trial data can be written as

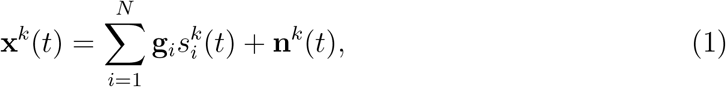

where **n**^*k*^(*t*) models the noise including the spatially correlated brain noise during the *k*-th epoch and is assumed uncorrelated between epochs, i.e. *E*{**n**^*k*^(*t*)**n**^*l*^(*t*)} = 0, *k* ≠ *l*, and 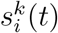 are source timeseries.

It is implicitly assumed that only a small fraction of *N* sources is active and the summation in (1) can be replaced by a smaller summation over a subset Ω of active sources. However, since the subset of active sources is unknown, we will stick to the more general formulation as presented above so that inactive sources are assumed to have *s_i_*(*t*) = 0 over the duration of the trial.

When estimating functional netwroks we are primarily interested in the coherent induced activity, i.e. rhythmic signals that demonstrate task specific (and locked to the stimulus) increase of functional coupling and in connectivity studies we typically exclude the evoked response from consideration. Therefore in the above we will assume that 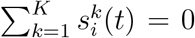 which is achieved in practice by subtracting the evoked response 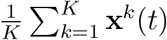 from each trial.

Networks analysis is traditionally performed via two steps. At first one uses an inverse solver to obtain estimates of the source activation profiles *s_i_*(*t*). Once this is done, within the second step the functional coupling metrics are computed to analyze the functional networks of interest.

To analyze within frequency linear synchronization we typically for each pair of source timeseries estimates *s_i_*(*t*) and *s_j_*(*t*) compute the coherence function *coh_ij_* defined as normalized squared cross-spectral coefficient and is a function of frequency, i.e.

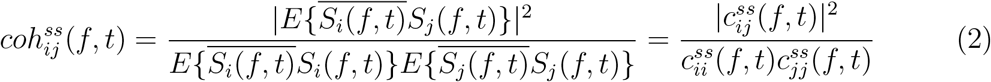

where *S_i_*(*f*, *t*) is the time-frequency representation of the *i*–*th* source timeseries and 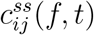 are the elements of the source space (note *ss* superscript) cross-spectral matrix 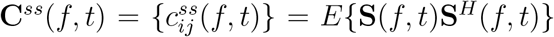 where *S*(*f*,*t*) = [*S*_1_(*f*,*t*),…,*S_N_*(*f*,*t*)]^*T*^ is a vector of time-frequency transformed source activation timeseries *s_i_*(*t*) for *i* = 1,…, *N*, *t* indexes the time within a trial and *f* is the frequency of interest. Averaging across *K* trials yields an estimate of the true coherence between source timeseries. In practice the actual source timeseries in (2) are replaced with their estimates and the expectation operation is replaced by averaging across trials for stimulus-based paradigms or across time within a sliding window for the resting-state data analysis. Both techniques allow us to retain the temporal dimension of coherence and subsequently analyze it as a time-varying function.

From (2) we can see that the source space cross-spectral matrix **C**^*ss*^(*f*,*t*) is the sufficient statistics for calculating the coherence for all pairs of sources and therefore is of interest. PSIICOS relies on the generative equation linking the source and sensor-space cross-spectral matrices which we derive next.

### 2.2. Generative model for the cross-spectrum

Applying a time-frequency transform (typically wavelet transform) to both sides of (1) and exploiting the transform’s linearity we will obtain the following expression for the time-frequency transformed sensor signals

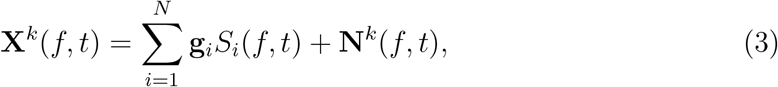

Then, by applying the definition of the cross-spectral matrix to the time-frequency transformed sensor data (3) we obtain the following generative model of the sensor space cross-spectrum **C**^*XX*^(*f*,*t*) linking it to the source space cross-spectral matrix **C**^*ss*^(*f*,*t*):

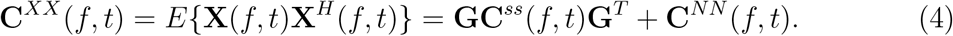

Here **C**^*NN*^(*f*,*t*) is the cross-spectral matrix of the additive noise and **C**^*ss*^(*f*,*t*) = *E*{**S**(*f*,*t*)**S**^*H*^(*f*,*t*)}. The approximation of the expectation operation *E*{} is computed by averaging the products of time-frequency coefficients matrices over *K* trials.

### 2.3. Manifestation of the SL effect in the cross-spectrum

Similarly to the way it was done in (Ossadtchi et al., 2018) we expand matrix products in (4) into the summations of the modulated outer products of topographies and split the real and imaginary parts to obtain:

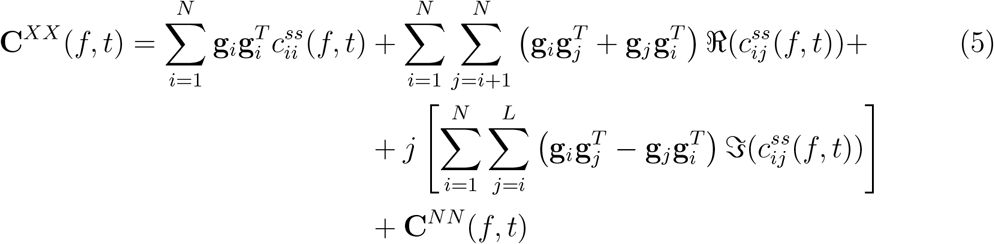

Let’s now consider the diagonal of the source space cross-spectrum matrix **C**^ss^(*f*, *t*) in more detail. The elements of the main diagonal of this matrix are real numbers and they represent the power of source activation timeseries around frequency *f* at time *t*. The structure of the generative model reflects the fact that after being mapped from the source space to the sensor space using the operator **G**, these power terms will manifest themselves both in the diagonal and off-diagonal elements of the sensor-space cross-spectral matrix **C**^xx^(*f*, *t*). The *i*-th row of matrix **G** corresponds to the lead field of the *i*-th sensor and describes how the activity of neuronal sources gets mixed in the sensor measurements.

The symmetric matrices 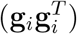 map the diagonal elements of the source-space cross-spectrum matrix (source powers) onto the sensor-space cross-spectrum, another set of symmetric matrices 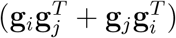 carries information about the *real* part of the off-diagonal source space cross-spectrum elements, while the anti-symmetric matrices 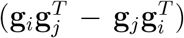 map the *imaginary* part of the off-diagonal elements, 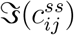. The orthogonality of symmetric and anti-symmetric matrices leads to the independence of the imaginary part of the sensor-space cross-spectrum on the spatial leakage. This observation led to the development of several SL-invariant functional coupling metrics (Nolte et al., 2004; Stam et al., 2007; Nolte et al., 2008; Pascual-Marqui et al., 2011; Ewald et al., 2012). These elegant solutions come at a price of neglecting the functional networks whose timeseries exhibit zero or close-to-zero mutual phase lag.

In (Ossadtchi et al., 2018) we introduced the PSIICOS framework based on the projection operation in the *M*^2^-dimensional space of the vectorized sensor space cross-spectral matrices, PSIICOS framework allows for assessing both real and imaginary parts of the source-space cross-spectrum(Ossadtchi et al., 2018). In the following sections, we will analytically demonstrate that this projection is optimal in terms of minimizing the amount of common mode signal in the estimates of activity for the target sources being assessed for connectivity.

### 2.4. Deriving the method of optimal filtering

It is important to note that the SL effect is not limited to the cross-talk only between the two sources for which the connectivity is being measured. Any source that leaks into the estimate of activity for both of these sources simultaneously will potentially result in erroneous estimates. Therefore, in order to accurately evaluate connectivity between a pair of sources, it is in general necessary to design a set of spatial filters that maximally suppress cross-talk from sources that are simultaneously leaking into the estimates of activity for both target sources.

To quantify the amount of such leakage we can use the concept of resolution kernel (RK) *R*(**r**, **r′**) introduced in the theory of inverse problems and used in (Sekihara and Nagarajan, 2008)). Let us consider an inverse operator that recovers the activity at a point **r** in the cortex based on measurements from sensors, and fix another cortical point **r′**. The function *R*(**r**, **r′**) indicates the amount of signal that leaks from point **r′** into the estimate of activity of a source located at **r**. For linear methods of solving the inverse problem, the function *R* can be expressed as a scalar product as follows:

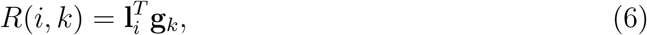

where **l**_*i*_ is the filter used to estimate the activity of the *i*-th cortical source and ***g**_k_* is the topography of the *k*-th source.

We can define a vector-valued function **B**(*i*), where *i* = 1,…,*N*, that quantifies the amount of signal flow from each cortical point to the fixed point *i*:

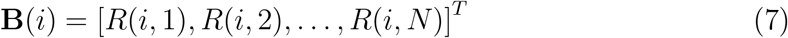

The function has been referred to as the Beam Response (BR) (Sekihara and Nagarajan, 2008) and is similar to the concept of a lead-field, which represents a row of the forward matrix that shows the sensitivity of a specific sensor to the activity of cortical sources. Vector **B**(*i*) reflects the amount of signal leaking from each cortical source into the estimate of activity of the *i*-th *source*. In the ideal case, the *i*–th element of **B**(*i*) is one, and the rest of its elements are zeros, i.e. **B**(*i*)_*n*_ = *δ*(*i* – *n*), for *n* =1,…,*N*.

Let us now analyze two points *i* and *j* on the cortex. The degree of overlap between vectors **B**(*i*) and **B**(*j*) plays a crucial role in determining the magnitude of the shared component of the SL signal. In an ideal scenario, if the components of both vectors, **B**(*i*) and **B**(*j*), are non-negative and orthogonal to each other, the common mode signal flowing simultaneously through points *i* and *j* will be null, resulting in the elimination of the SL and its contribution to the connectivity estimate. This highlights the importance of designing a set of filters that minimize the overlap between **B**(*i*) and **B**(*j*) for every pair of points.

In order to construct such filters, it is essential to establish a methodology for measuring the overlap between the vectors **B**(*i*) and **B**(*j*). The scalar product of these vectors may initially seem like an adequate measure of the overlap: if this measure is equal to zero, the common leakage is absent. However, this measure is flawed due to the potential sign change of the components of these vectors. Even though the scalar product between vectors **B**(*i*) and **B**(*j*) is zero, indicating that they are orthogonal, the individual components of these vectors may still be substantial and interfere with each other if they possess opposite signs. For instance, if the functions **B** are equal to (1, −1) and (1, 1) correspondingly for the points *i* and *j*, the BRs are orthogonal to each other, but the SL between the sources is significant.

To resolve this issue, a more accurate measure of overlap is required. The magnitude of the scalar product of the vectors **B**(*i*) and **B**(*j*) whose components are first squared, serves as a more appropriate measure of overlap. We will refer to the result of this scalar product, the function *μ*(*i*,*j*), as *mutual spatial leakage* (*mutual SL*) of the points *i* and *j* and formally define it as:

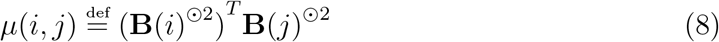

where (·)^⊙2^ notation is inspired by the Hadamard product and denotes squaring the corresponding vector element-wise. This measure is similar to that used in (Nolte et al., 2009) for describing the constraints on the theoretical distribution of sources.

Taking into account the definition of resolution kernel (6), the mutual SL can be expanded as:

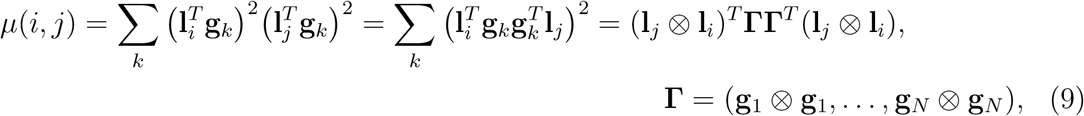

note that **g**_*j*_ ⊗ **g**_*i*_ is the Kronecker product based notation equivalent to 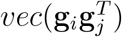 used in the original PSIICOS paper (Ossadtchi et al., 2018).

With the aim of minimizing mutual SL while preserving the signal of interest, we seek to find a pair of filters **l**_*i*_, **l**_*j*_ for points *i* and *j*. To achieve this, it is necessary to impose additional constraints on the length and orientation of these spatial filters. One possible constraint is the requirement that the signal amplification coefficient for signals originating from points with topographies **g**_*i*_ and **g**_*j*_ is equal to 1. Taking into account this constraint, the optimization problem for finding the pair of filters **l**_*i*_, **l**_*j*_ can be written as follows:

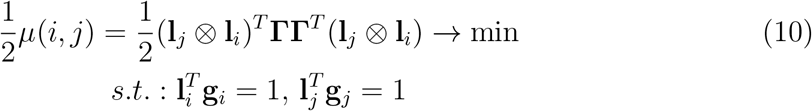

Under this constraint, the optimization problem cannot be explicitly solved by the method of Lagrange multipliers due to the presence of the Kronecker product in the objective function. Thus, it must be solved numerically, for example, by the gradient descent method. However, this problem can be significantly simplified generalizing the concept of filters to the *M*^2^-dimensional space of the vectorized MEG or EEG data covariance matrices.

It should be noted that the constraints of (2.4) imply that

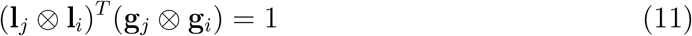

Note that the constraint, written in such a form as the objective function in equation 2.4, depends on the filters **l**_*i*_, **l**_*j*_ only through their Kronecker product, which we denote as **v**_*ij*_. For simplicity, we also denote **g**_*j*_ ⊗ **g**_*i*_ as **q**_*ij*_. We will refer to such *M*^2^-dimensional “topography” of a dyadic network as 2-topography.

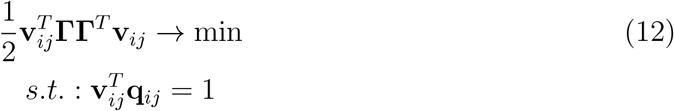

Such a problem regarding the variable **v**_*ij*_ can be easily solved using the Lagrange multiplier method. Its Lagrangian can be written as follows:

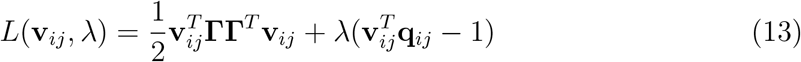

Differentiating the Lagrangian with respect to **v**_*ij*_, we obtain:

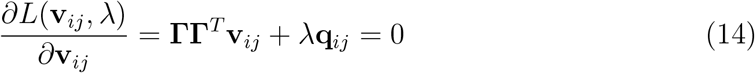

If λ ≠ 0, the equation 14 has no solutions: the columns of the matrix **Γ** are vectorizations of symmetric matrices, and thus the product **ΓΓ**^*T*^**v**_ij_ is also a vectorization of a symmetric matrix; however, **q**_*ij*_ is not a vectorization of a symmetric matrix. The difference is in our case for λ ≠ 0 we have an incongruent system of equations due to the described above special structure of **Γ** matrix and vector **q**_*ij*_. This means that we need to set λ = 0 and obtain the following equation for **v**_*ij*_:

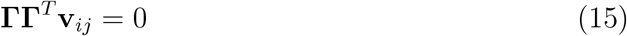

It follows that the vector **v**_*ij*_ must belong to the orthogonal complement of *col* **Γ**, the column space of **Γ**. This means that the vector **v**_*ij*_ can be obtained as the result of applying the projection **P** = **I** – **ΓΓ**^†^ to some vector **ṽ**_*ij*_ from the general linear space of dimension *M*^2^.

Differentiating the Lagrangian in equation (13) with respect to λ results in:

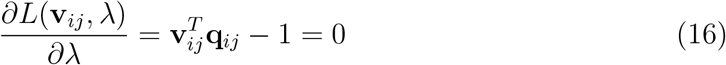

Taking into account the general form of the vector **v**_*ij*_, as well as the idempotency and symmetry of the projection operator, we will obtain:

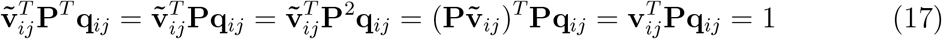

The last equation defines a hyperplane in the affine subspace of vectors orthogonal to the columns of the matrix **Γ**. As a result, the optimization problem has an infinite number of solutions. To choose a unique solution, we can use the criterion of minimal norm as previously. Among the vectors of this hyperplane, the one with the minimum norm satisfying (17) has to be the vector collinear to **Pq**_*ij*_:

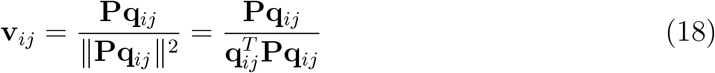

Therefore, the filter that minimizes mutual SL for cortical points *i*, *j* is obtained as a result of the projection **P** applied to the corresponding 2-topography. Namely this projection was previously described and evaluated in Ossadtchi et al. (2018) where the authors basically suggest first to project the vectrorized cross-spectrum *vec* (**C**^XX^(*f*, *t*)) using operator **P** and then perfrom a scan using **Pq**_*ij*_ vectors, see Figure 1 in (Ossadtchi et al., 2018). Due to idempotency of **P** this is equivalent to using filter (18).

**Figure 1:**
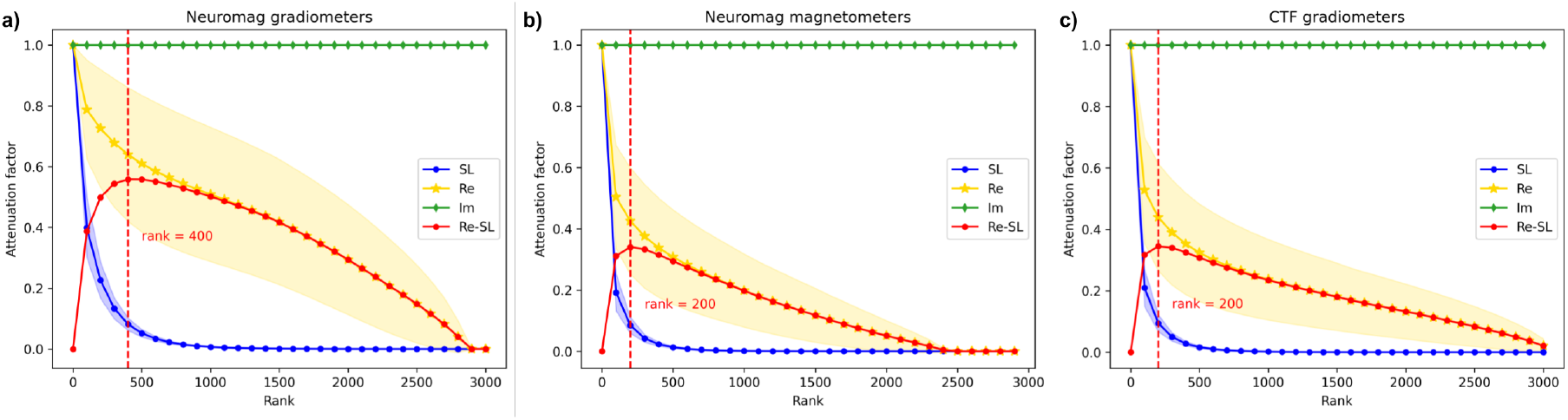
SL and Re subspace power suppression and the suppression difference curve for the original (projected) PSIICOS as a function of projection rank a) for Neuromag 204 planar gradiometers probe, b) for Neuromag 102 megnetometers probe and c) for 275 radial gradiometers CTF probe

In Ossadtchi et al. (2018) the authors applied PSIICOS to estimating real and imaginary part of the source-space cross-spectrum separately which leads to the normalization imbalance when using the real and imaginary parts simultaneously. To address this issue, we modify the constraint in (2.4) to read as 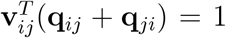 and obtain the optimal filter to recover the real-part of the sources space cross-spectral coefficient 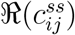:

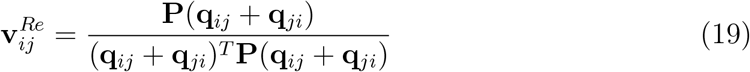

For the imaginary part the constraint is 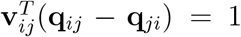 which leads to the following filter:

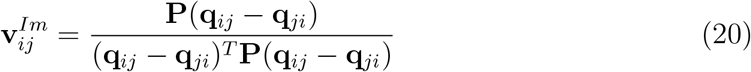

Since (**q**_*ij*_ – **q**_*ji*_) and the column space of **Γ** are orthogonal, (20) can be restated as

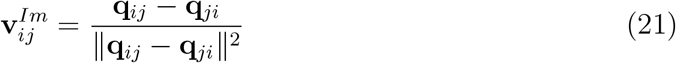

Note that in the limiting case when the number of cortical nodes exceeds the value *M*(*M* + 1)/2, equal to the total number of symmetric *M* × *M* matrices, the application of operator **P** completely nullifies the vectorized symmetric matrices. This is effectively equivalent to taking the imaginary part of the cross-spectrum, as the imaginary parts of the antisymmetric topographies are antisymmetric and remain unchanged under projection.

In practice we operate with MEG or EEG timeseries data projected into a lower dimensional principal subspace of the forward model matrix **G**. Typically, for EEG and MEG the rank of this subspace falls in 60 to 150 range. In MEG, when dealing with MaxFilter processed planar magnetometer data, the resulting rank is typically chosen to be below 100. If we consider a 73× 73 matrix that is symmetric, the total number of such matrices is given by *n* = 73 × (73 + 1)/2 = 2701. Therefore, even for a relatively sparse cortical mesh with greater than *n* vertices, application of the full rank **P** to the sensor-space cross-spectrum will eliminate any information about the real-valued component of the cross-spectrum, see Figure 1. b.

To balance the trade-off between the SL suppression and retention of the information about the real components of the source-space cross-spectrum we limit the projection rank *R*. Rank-limited **P**^*R*^ is computed using the first *R* left singular vectors **u**_*i*_, *i* = 1,…, *R* of **Γ** corresponding to the first *R* largest singular values as

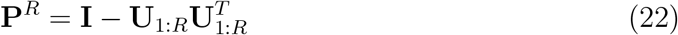

where **U**_1:*R*_ = [**u**_1_,**u**_2_,…**u**_*R*_]

In Figure 1 we show the average suppression the PSIICOS projection bestows upon the spatial leakage (SL), real (Re) and imaginary (Im) exemplar 2-topographies for different projection ranks *R*. To obtain these plots we performed 10000 Monte-Carlo (MC) trials. At each we randomly chose vertex index pairs (*i*, *j*) and computed the ratio of the post-to pre-projection norms of the corresponding 2-topographies **g**_*i*_ ⊗ **g**_*i*_ + **g**_*j*_ ⊗ **g**_*j*_, **g**_*i*_ ⊗ **g**_*j*_ + **g**_*j*_ ⊗ **g**_*i*_ and **g**_*i*_ ⊗ **g**_*j*_ − **g**_*j*_ ⊗ **g**_*i*_. As expected, the 2-topographies corresponding to the imaginary part of true source space coherence appeared to be not affected by the PSIICOS projection. For the 2-topographies corresponding to the SL component the suppression rapidly grows with the rank increase while Re components are only reasonably suppressed. As evident from the suppression difference curve (SL-Re), rank values *R^NMG^* = 400 and *R^CTF^* = 200 result into the best trade-off between the SL and Re components suppression and therefore can be chosen as optimal. Theoretically, the projection rank choice may also be affected by the fact that due to the inherent properties of the electromagnetic forward problem the numerically stable rank of col(**Γ**) is typically less than the number of sources (number of columns of **Γ**). However, based on our experience the numerical rank of **Γ** is usually higher than *R^NMG^* = 400 and *R^CTF^* = 200 for the Neuromag and CTF probes correspondingly.

### 2.5. Regularized PSIICOS solution

As already noted, with a sufficiently large number of cortical points, **P** constructed without the projection rank restriction completely eliminates information about the real parts of the source space cross-spectral matrix off-diagonal elements. Limiting the projection rank allowed us to manipulate the trade-off between the SL suppression and Re components power retention. In the previous derivation we have chosen the unique solution using the standard minimum-norm argument, see (18).

Here we consider an alternative way of formulating and solving the PSIICOS optimization problem. The approach is based on the Tikhonov regularization and allows us to simultaneously balance between the minimum norm and minimal leakage requirements. The optimization problem in this case is written as

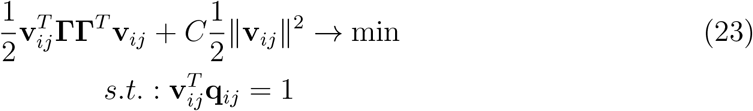

where 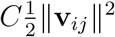 allows for a soft control over the norm of the obtained PSIICOS filter vector.

The Lagrangian for this optimization problem is

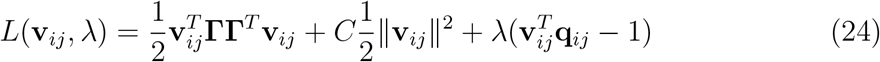

Taking the derivative with respect to **v**_*ij*_ we get:

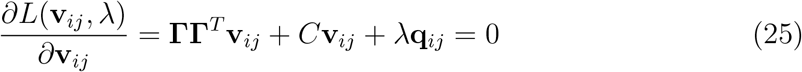

This time, thanks to the regularization (25) has solutions for non-zero λ

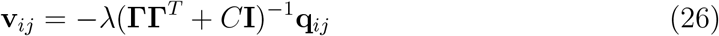

Multiplying the left and the right parts by **q**_*ij*_ and taking into account the constraint 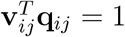, we get the expression for λ:

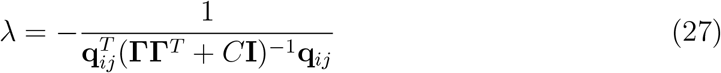

The corresponding filter **v**_*ij*_ is then given by

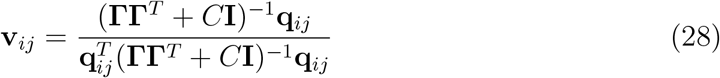

As it turns out both (18) and (28) give nearly identical results for a broad range of regularization parameter *C*, compare Figures 1 and 2.

**Figure 2:**
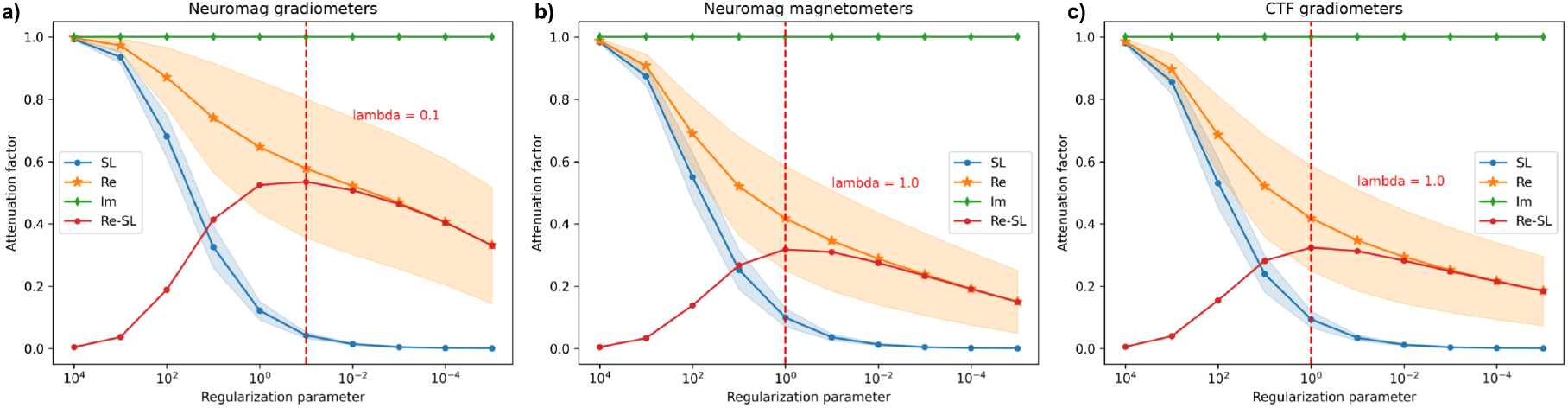
SL and Re subspace power suppression and the suppression difference curve for the regularized PSIICOS as a function of the regularization parameter a) for Neuromag 204 planar gradiometers probe, b) for Neuromag 102 megnetometers probe and c) for 275 radial gradiometers CTF probe

It is also possible to reformulate the constraint in (2.5) to specifically target the real and the imaginary parts of the the source space cross-spectral coefficient *c_ij_* between the timeseries of the *i*–th and *j*–th sources, see (5).

The optimal filter for the real part of the *ij*–th source-space cross-spectral coefficient results from imposing 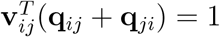 constraint:

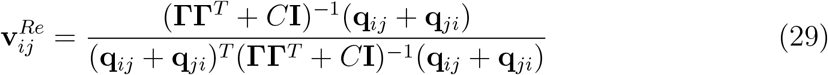

Correspondingly, for the imaginary part of the *ij*–th source-space cross-spectral coefficient with 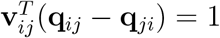 constraint we get:

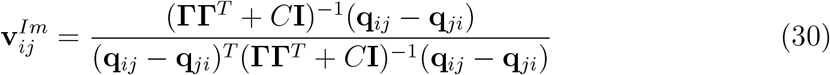

Note that due to the orthogonality of (**q**_*ij*_ – **q**_*ji*_) and the column space of the spatial leakage matrix **ΓΓ**^*T*^, equation (30) can be restated as

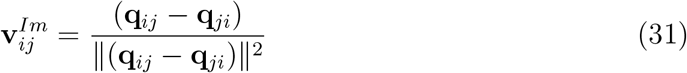

Interestingly, filter (31) is simply a match filter in the *M*^2^-dimensional space and tuned to the 2-topography of the network with nodes in the *i*-th and *j*-th vertices of the cortical mesh. Theoretically speaking, 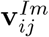 is a maximum-likelihood (ML) estimator of 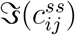 in the case when the observation noise is spatially white whose covariance is identity and vanishes from the corresponding expression for the ML estimator. Filter 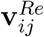 can also be viewed as an ML estimator of 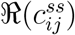 where the noise comes from the spatial leakage with *M*^2^ × *M*^2^ “spatial” covariance **ΓΓ**^T^ and the additive noise term λ**I**. Clearly, these estimators of network activity can be extended to include the information about the activity of other discrete networks, as it is done in a more traditional ML estimation of neuronal sources nicely summarized in (Hauk and Stenroos, 2014). For this, the set of constraints (2.4) and (2.5) needs to be augmented to include the explicit 2-topographies of the potentially interfering networks, see *Discussion* section.

Regularization parameter *C* in (2.5) serves the role similar to that of the rank parameter *R* in the original PSIICOS approach. Adjusting *C* we can softly tune the thresholding and exclude from the inversion process matrix components corresponding to low eigenvalues. Projection with rank *R* and the regularization based scheme can be put in correspondence by choosing for example regularization parameter *C* equal to the *R*-th singular value *σ_R_* of **ΓΓ**^*T*^ so that the corresponding eigenvalue of *C*(**ΓΓ**^T^ + *C***I**)^-1^ is equal to *C*/(*C* + *σ_R_*) = *σ_R_*/(*σ_R_* + *σ_R_*) = 0.5.

The suppression factor plots sampled at the corresponding (*R*, *C*) pairs for the projected and regularized PSIICOS are shown in Figure 3.a - c. for the Neuromag planar gradiometers, magnetometers and CTF radial gradiometers correspondingly. As expected the two approaches appear to be nearly equivalent. The suppression factor difference plots presented in panels c - f allow for selection of the optimal values controlling the regularization process in both approaches.

**Figure 3:**
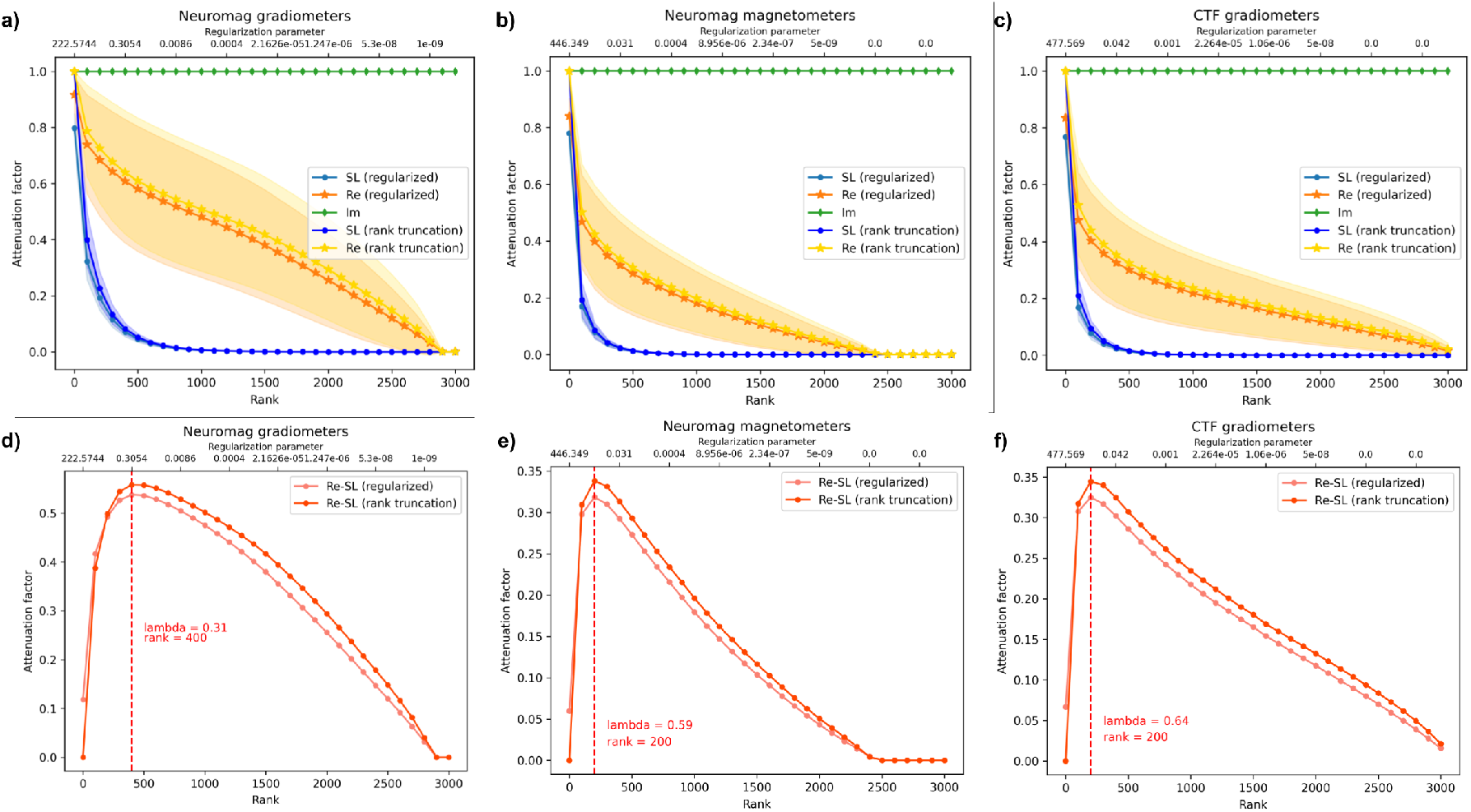
SL and Re components power suppression curves a),b),c), and the suppression difference curves c),d),f) as functions of projection rank *R* and regularization parameter *C*. The curves are sampled at the corresponding values of the argument by choosing *C* = *σ_R_* for 204 Neuromag planar gradiometers probe (a,d), 102 Neuromag magnetometers probe (b, e) and 275 radial gradiometers CTF probe (c,f)

## 3. Lower rank approximation

The vast majority of methods for the source-space functional coupling assessment is based on precomputing the source timeseires using an inverse solver of choice followed by computing the statistics of interest. For example, the DICS (Groß et al., 2001) technique is based on the analysis of the estimate of the source space cross-spectral coefficient *c_ij_* obtained as

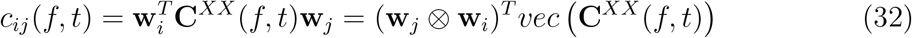

where **w**_*i*_ and **W**_*j*_ are the spatial filters tuned for the *i*-th and *j*-th source correspondingly. The operation described by (32) can be restated as

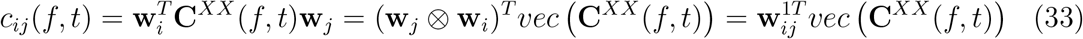

Vector 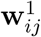 represents a filter operating in the *M*^2^ dimensional space and acting on the vectorized cross-spectrum *vec* (**C**^XX^(*f*, *t*)). In the traditional connectivity estimation setting (32) this filter has a special constraint that forces it to be expressed as a Kronecker product of two *M*-dimensional vectors, e.g. 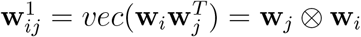. In what follows this particular structure will be denoted as the rank-1 form.

As demonstrated in the previous sections the PSIICOS approach relaxes this constraint on the spatial filter structure and seeks for a *M*^2^ × 1 **w**_*ij*_ to optimise the trade-off between the extent of the mutual leakage suppression and the amount of the retained power modulated by the real component of the source-space cross-spectrum. The latter is achieved either by adjusting the projection rank *R* or the regularization parameter *C* whose conceptual equivalence was demonstrated in Figure 3. Operation in the *M*^2^ dimensional space endows PSIICOS with extra power in suppressing the mutual spatial leakage component as compared to the traditional processing described by equation (32). It allows PSIICOS to efficiently attenuate the mutual spatial leakage (8) in a principled way and retain information about the sensor-space cross-spectral components modulated by the real part of the source-space cross-spectrum.

How much fidelity in suppressing the SL will is going to be lost if we approximate the PSIICOS filter using rank-1 representation? To answer this question we first need to come up with approximation quality criterion *Q*. One possible approach is to gauge the approximation quality using the norm of the difference between the original **w**_*ij*_ and the rank-1 approximated PSIICOS filter 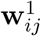. We dub this approach as “naive” for the reasons that will become clear later, 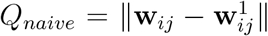. Optimizing *Q_naive_* leads to the following algorithm for building a rank-1 approximation of 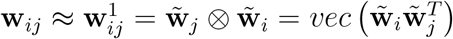:

- Reshape PSIICOS **w**_*ij*_ into a *M* × *M* matrix **W**_*ij*_
- In order to find such 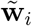 and 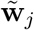 that 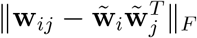 is minimum use the defining property of the SVD and represent **W**_*ij*_ as **W**_*ij*_ = **USV**^*T*^, with **U** = [**u**_1_,**u**_2_,…, **u**_*M*_], **V** = [**v**_1_**v**_2_,…,**v**_*M*_], **S** = **diag**([*s*_1_,*s*_2_,…,*s_M_*])
- Compute rank-1 approximation of **w**_*ij*_ as 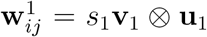 as the vectorized outer product of the first pair of singular vectors corresponding to the largest singular value *s*_1_, and assign 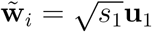 and 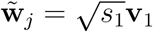
- Higher rank approximations can also be computed as 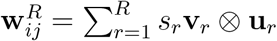

Per defining property of SVD the above approach minimizes the residual difference 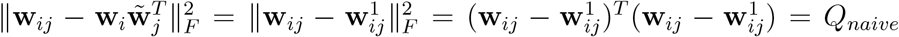 which does not control the extent to which the mutual spatial leakage *μ*(*i*, *j*) is suppressed. The above approximation scheme can be improved to take this into account in order to obtain a better approximation. To control *μ*(*i*, *j*) and achieve the best low rank approximation of a filter in a sense of retaining maximum of the SL attenuation we will reformulate our approximation quality criterion as

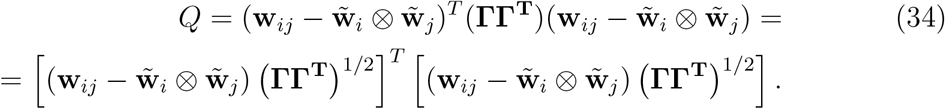

*Q* in (34) can be optimized in much the same way as *Q_naive_* except that prior to reshaping and performing the SVD the original filter **w**_*ij*_ needs to be multiplied by the scaling matrix **H** = (**ΓΓ**^T^)^1/2^. After the SVD, the obtained lower rank approximation needs to be reshaped and multiplied back by the inverse of **H**.

The left and the right panels of Figure 4 compare the two lower rank approximation methods by visualizing the extent to which the spatial leakage (SL) subspace variance projects onto the specific filter designed to recover source space cross-spectral coefficient *c_ij_* for a randomly chosen pair of cortical vertices (*i*,*j*). The ordinate of the dots in this plot corresponds to the amount of mutual leakage 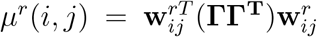 for *r* = 1, 2, 50 and for the full rank and measures the amount of average residual spatial leakage power after applying the PSIICOS filter. The abscissa of each dot reflects the correlation coefficient of the topographies of sources located in the *i*-th and *j*-th vertex. We have also normalized each approximation to ensure the unity gain constraint in the ‘‘direction” of the 2-topography **g**_*j*_ ⊗ **g**_*i*_ + **g**_*i*_ ⊗ **g**_*j*_ modulated by the real part of the source space cross-spectrum *c_ij_* coefficient. As we can see the second scheme (on the left) achieves smaller *μ^r^*(*i*,*j*) for all approximation ranks *r* as compared to the naive approach (on the right).

**Figure 4:**
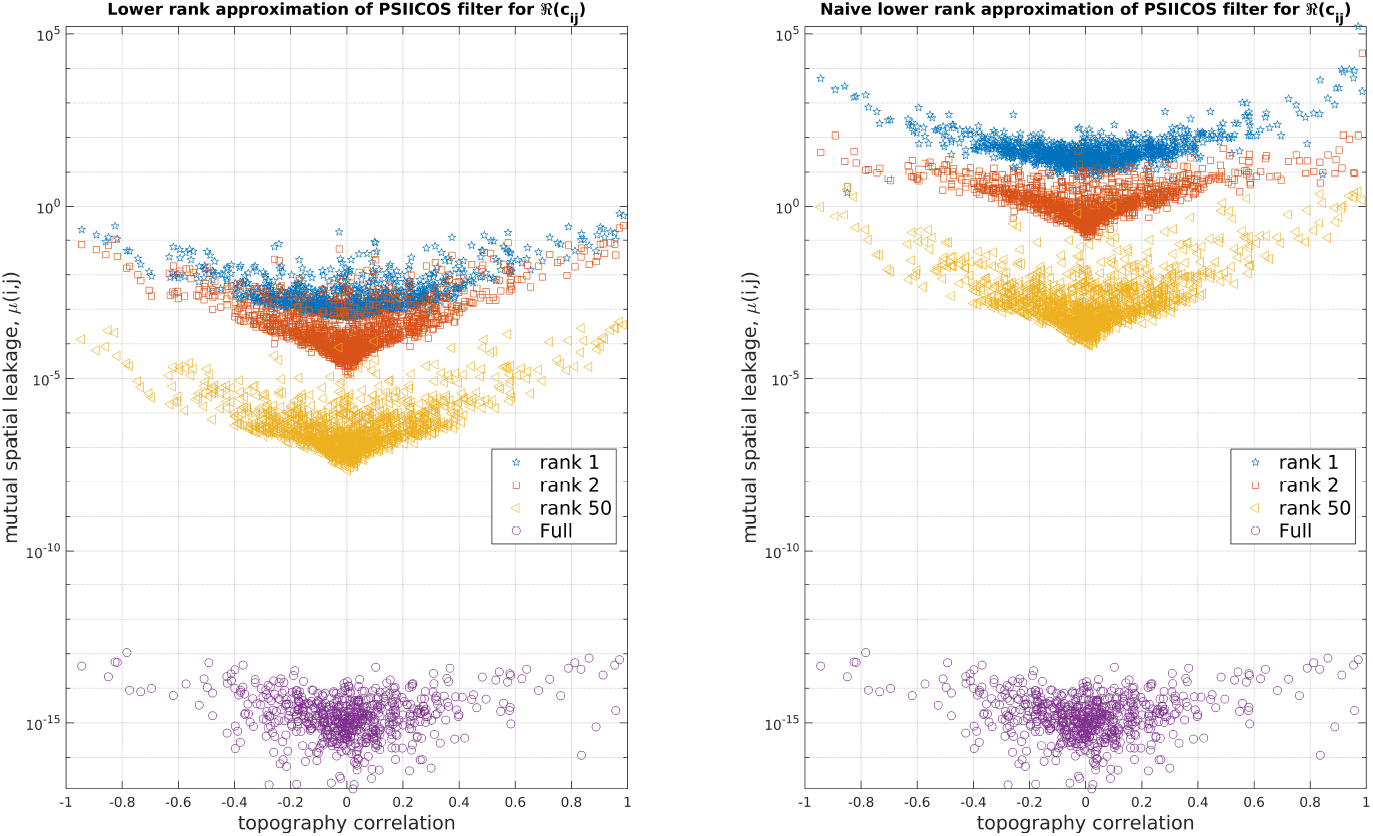
Naive vs. optimal lower rank approximation of PSIICOS filters. The ordinate of the dots in this plot corresponds to the mutual spatial leakage *μ^r^*(*i*,*j*) computed as 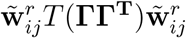 for different approximation ranks *r* = 1, 2, 50 and for the full rank. We have also normalized each approximation to ensure the unity gain constraint in the ‘direction” of the 2-topography **g**_*j*_ ⊗ **g**_*i*_ + **g**_*i*_ ⊗ **g**_*j*_ of the real part of the source space cross-spectral *c_ij_* coefficient. Each dot corresponds to a randomly chosen pair (*i*,*j*) of cortical sources. Dot’s position along the horizontal axis is determined by the correlation coefficient of the *i*–th and the *j*–th source topographies 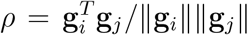. As we can see the optimal scheme (on the left) allows for a stronger attenuation of the mutual spatial leakage as compared to the naive scheme (on the right).

In (Ossadtchi et al., 2018) the authors suggested conducting a reduced version of the MUSIC scan to identify active networks. Since the MUSIC family of techniques is highly sensitive to the forward model accuracy when evaluating the quality of the approximation, we must consider the forward model similarity metric alongside the mutual spatial leagake suppression criterion. PSIICOS filters **v**_*ij*_ are basically match filters tuned to the SL-debiased 2-topography and we will measure the cosine of the angle between the approximated and the original filter as 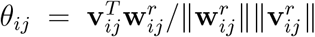. Figure 5.a shows the distribution of *θ_ij_* for a set of randomly selected pairs (*i*,*j*) of cortical locations. As in Figure 4.a the horizontal coordinate corresponds to the *i*-th and *j*-th topography correlation coefficient 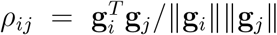. We can see that the increase of the approximation rank reduces the fraction of examples with low cosine similarity. Typically, locating cortical sources using the MUSIC technique requires subspace correlation values above 0.95, which means that the obtained rank-1 approximation is likely not sufficient to reliably identify networks whose activity is manifested in the observed sensor-space cross-spectrum.

**Figure 5:**
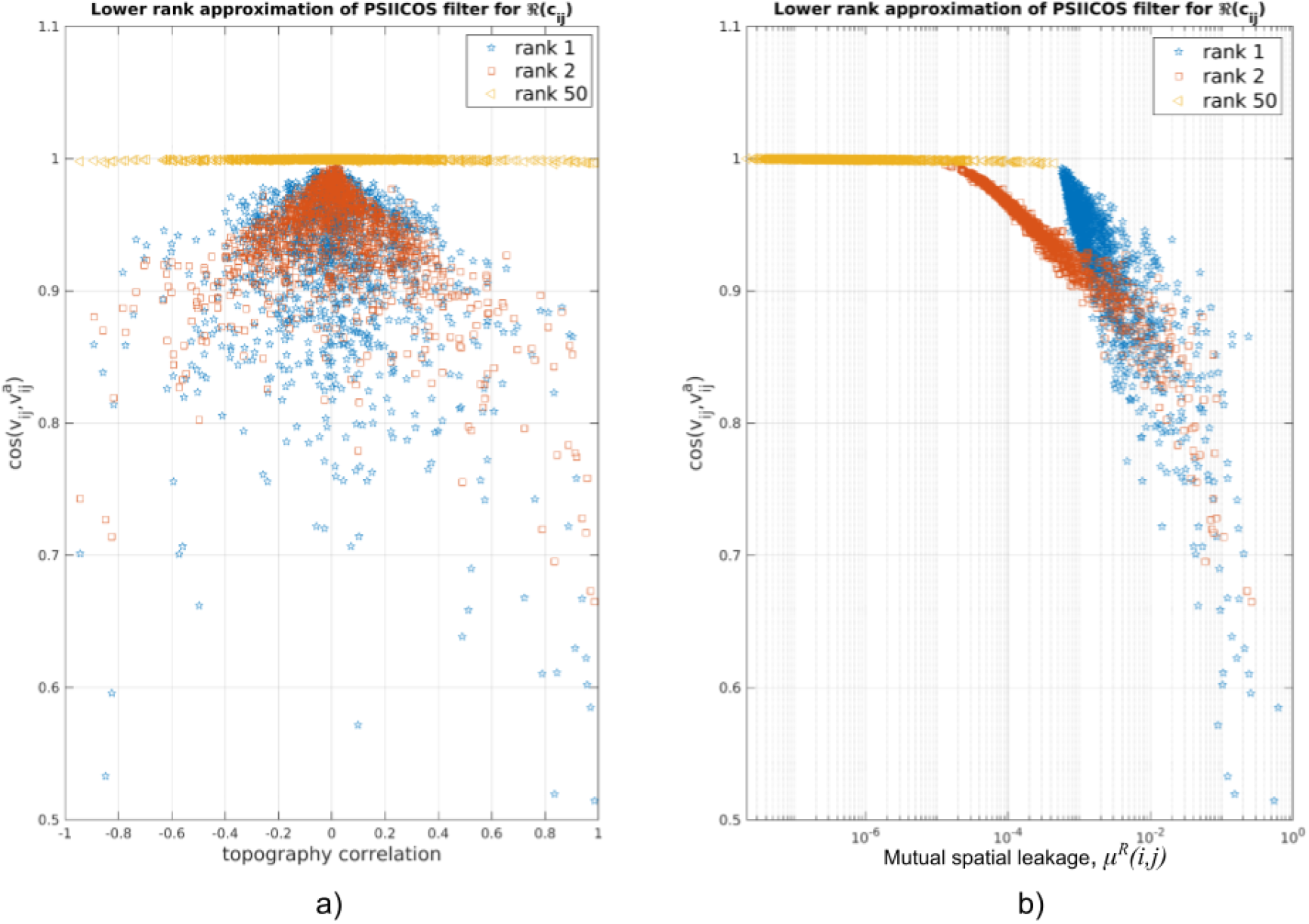
Lower rank approximation quality assessment. a) The distribution of *r_ij_* for a set of randomly selected cortical location pairs (*i*, *j*). The horizontal coordinate corresponds to the *i*-th and *j*-th topography correlation coefficient *ρ*. The increase of the approximation rank reduces the fraction of examples with low cosine similarity. b) Cosine similarity vs SL attenuation for a set of randomly chosen cortical location pairs (*i*, *j*). The increase of the approximation rank orients the distribution of points more horizontally and shifts it to the left.

Figure 5.b provides an easy-to-understand summary of our results for this section. Each dot in the graph corresponds to a randomly chosen pair of cortical locations (*i*, *j*). The mutual spatial leakage *μ^r^*(*i*, *j*) and cosine similarity criteria are used as coordinates for the dots. As the rank of the approximation increases, the distribution of dots becomes more horizontal around 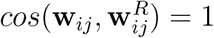 and shifts towards the left indicating average mutual spatial leakage reduction.

## 4. Discussion

We have demonstrated that the PSIICOS projection is optimal in a sense of minimizing the overlap between beam response vectors corresponding to the cortical locations evaluated for functional connectivity. We defined a new metric of overlap, which we referred to as “mutual spatial leakage,” in equation (8). This metric is similar to that used by (Nolte et al., 2009; Marzetti et al., 2008) to reflect the properties of the hypothesized underlying ground truth distribution of neuronal sources.

In the extreme case PSIICOS projection leads to a complete depletion of mutual spatial leakage power, resulting in the established approaches that rely solely on the imaginary part of the sensor-space cross-spectrum (Nolte et al., 2004). However, these techniques fail to detect functional networks with nodes coupled at or near zero phase lags since this information is stored in the real part of the cross-spectrum.

PSIICOS operates in the *M*^2^ dimensional product space of sensor signals, enabling direct assessment of the spatial structure of mutual spatial leakage and identifying a set of complementary subspaces capturing the largest mutual spatial leakage variance for a given subspace rank. This approach facilitates tuning the mutual spatial leakage subspace rank to balance between suppressing mutual spatial leakage and retaining information about the real part of the source-space cross-spectrum.

Additionally, we have demonstrated an alternative way to arrive at PSIICOS which relies on using Tikhonov regularization. This approach produces a solution that is very similar to the PSIICOS projection, but instead of controlling the retention-suppression trade-off through the projection rank, it is controlled by the regularization parameter. The regularized PSIICOS approach offers a more conventional interpretation of the PSIICOS framework, which is parallel to the terminology used in EEG and MEG based source estimation literature. Instead of the concept of a neuronal source approximated by a single equivalent current dipole (ECD) in the conventional inverse problem framework, the PSIICOS paradigm introduces the notion of a dyadic network whose nodes are two ECDs located at disjoint locations of the cortex. Instead of conventional source topography, the PSIICOS operating with dyadic networks uses 2-topography computed using the Kronecker product of conventional ECD topographies of sources located in the corresponding nodes of the cortical mesh forming the dyadic network. Furthermore, instead of source activation timeseries, the PSIICOS framework employs cross-spectral coefficient time series *c_i,j_*(*t*, *f*), which characterizes the functional coupling within the elementary network formed by the *i*-th and the *j*-th cortical source.

In this light the solutions (29) and (31) are maximum likelihood (ML) estimators of the real and imaginary parts of the *c_ij_*, see equation (16) in (Hauk and Stenroos, 2014). Noise covariance matrix in (29) is formed by the mutual spatial leakage matrix **ΓΓ**^*T*^ and the added white noise term C**I** used for regularization. As an ML estimator, the algorithm basically performs match filtering in the space whitened by the inverse square root of **C**_*N*_ = **ΓΓ**^*T*^ + *C***I** noise covariance matrix. Solution (31) also represents a match filter tuned to the 2-topography **q**_*ij*_ – **q**_*ji*_ whose contribution to the sensor-space cross-spectrum is modulated by the imaginary part of *c_ij_*. In this case the noise covariance **C**_*N*_ is the identity matrix, i.e. **C**_*N*_ = **I**, which reflects the fact that the spatial leakage does not affect the imaginary part of the cross-spectrum. In fact, the estimates of 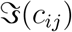 delivered by PSIICOS match those obtained with a more conventional imaginary DICS approach (Nolte et al., 2004; Ossadtchi et al., 2018).

Neither PSIICOS as described in (Ossadtchi et al., 2018) nor other functional connectivity methods (Bastos and Schoffelen, 2016; Greenblatt et al., 2012) explicitly account for the presence of multiple simultaneously active networks. The described link to the conventional ML estimators allows us to formulate a solution to the problem of estimating the activity of several concurrenty active networks by creating the appropriate constraints matrices 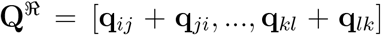 and 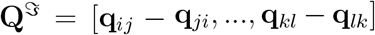 whose columns are the 2-topographies of the found to be active dyadic networks. Then, the matrix of the optimal in the ML sense filters for estimation the cross-spectral coefficients characterizing the activity of the active *c_ij_* will look like

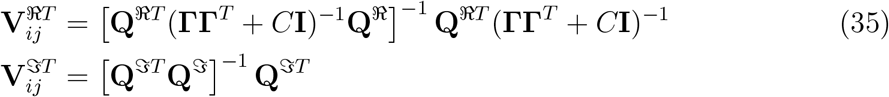

In the above equations the estimator is informed about the presence of active networks and will attempt to minimize the cross-talk in the estimates of the cross-spectral coefficients pertaining to these different networks. It also takes into account the noise that is primarily due to the mutual spatial leakage.

In a more realistic case when there is no information about the location of active networks and our goal is to find them, we can formulate a scanning beamformer like solution, (Sekihara and Nagarajan, 2008; Greenblatt et al., 2005). Beamformers can be both adaptive and non-adaptive. The non-adaptive beamformer will model the data covariance matrix using the forward model and assuming some statistical description of source activity. The detailed analysis of non-adaptive spatial filters in application to the conventioanl source estimation task is presented in (Sekihara and Nagarajan, 2008) and (Greenblatt et al., 2005). The key difference between the ML and the beamforming approaches lies in the specification of covariance matrix. The beamforming approach typically operates using the measurement covariance matrix and not only noise covariance as it is the case with the ML approach. Otherwise, the form of the solution remains the same as that obtained in the ML approach.

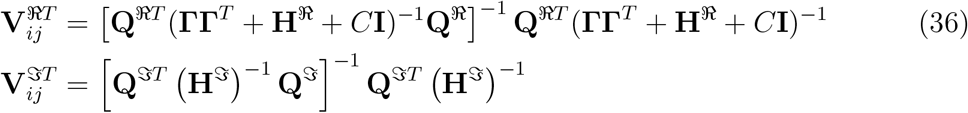

where matrices 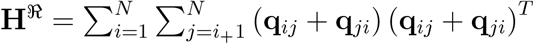 and 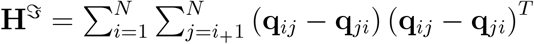 are the *M*^2^ × *M*^2^ “spatial” covariance matrices of dyadic networks activity as observed in the vectorized real and imaginary parts of the sensor space cross-spectrum.

To save on the degrees of freedom, we can also formulate a solution which would lie half-way between the non-adaptive spatial filter tuned to the dyadic networks whose 2-topographies are listed in **Q** matrices and a fully adaptive spatial filter solution. To this end we replace in (36) the product **ΓΓ**^*T*^ with 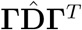 where 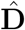 is a diagonal matrix whose elements 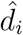, *i* = 1,…, *N* represent conventional source power estimates obtained with some other inverse solver, for example MNE. Also matrices 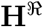 and 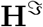 need to be reformulated as 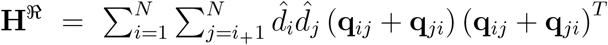 and 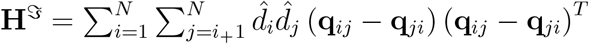 are the *M*^2^ × *M*^2^

It is also possible to formulate various solvers by working with the SL debiased cross-spectrum *vec*(**C**^*XX*^)^⊥*SL*^(*t*, *f*)

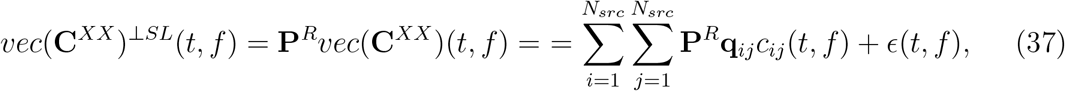

In this case, formally, the problem is identical to that of the classical source estimation except that instead of the ECD topographies as in (1) we are dealing with dyadic networks and their 2-topographies **P**^*R*^**q**_*ij*_ projected away from the principal spatial leakage subspace. In this setting the potential solvers for *c_ij_* (*t*, *f*) can be derived using Bayesian framework. The prior distribution of *c_ij_* (*t*, *f*) can be constructed using structural information obtained from the diffusion tensor imaging. It is also potentially possible to encode into complex-valued prior the expected neural pulse propagation speed which is known to depend on the myelinated fiber thickness (Schmidt and Knösche, 2019).

The spatial sparsity of the long range connections is explained by the small-worldedness property of the brain’s neural networks. Millimeter scale temporal resolution combined with characteristic times of binding in functional networks leads to temporal smoothness of the observed network activation profiles *c_ij_* (*t*, *f*). To model these two facts in the future we could use mixed norm priors (Gramfort et al., 2012).

The described potential benefits of PSIICOS stem from the linearity of the observation equation in the proposed formulations. The estimation task is then centered around the off-diagonal elements *c_ij_* of the source space cross-spectrum and not the normalized coherence values. This setting requires the development of the statistical testing methods that can be used to eliminate spurious detections, see (Kleeva and Ossadtchi, 2021) for the preliminary description of a possible approach.

PSIICOS computational requirements are modest. Creating the projection matrix **P** for a forward model with 1500 cortical vertices takes around 7 seconds on a regular PC and needs to be done only once for a given forward model. The vectorized scan over 1500 × 1500 pairs takes on the order of 0.5 seconds on the same machine.

Overall, the PSIICOS approach proposes a shift from the traditional two-step functional connectivity estimation procedure to operating in the M^2^-dimensional space of sensor-space cross-spectrum. This view allows us to highlight the fact that the classical two-step approach of connectivity estimation is simply a match filter that focuses on the topographies of the corresponding networks without considering the interference structure and therefore can be improved. PSIICOS implements only the most straightforward improvement and relaxes the requirement on the rank-1 structure of the filter. The future endeavors include exploring the efficiency of the formulated above novel estimators that follow well established set of principled assumptions and constraints and rest on the rich intuition accumulated in the community with respect to the traditional source estimation tasks (Hauk and Stenroos, 2014).

